# Electrocortical temporal complexity during wakefulness and sleep: an updated account

**DOI:** 10.1101/2020.02.20.958462

**Authors:** Joaquín González, Matias Cavelli, Alejandra Mondino, Claudia Pascovich, Santiago Castro-Zaballa, Nicolás Rubido, Pablo Torterolo

## Abstract

The states of sleep and wakefulness are critical physiological processes associated with different brain patterns of activity. The intracranial electroencephalogram allows us to measure these changes, thus, it is a critical tool for its study. Recently, we showed that the electrocortical temporal complexity decreased from wakefulness to sleep. Nevertheless, the origin of this complex activity remains a controversial topic due to the existence of possible artifacts contaminating the brain signals. In this work, we showed that complexity decreases during sleep, independently of the electrode configuration employed. This fact strongly suggests that the basis for the behavioral-state differences in complexity does not have an extracranial origin; i.e., generated from the brain.

## Introduction

The sleep-wake cycle is a critical physiological process and one of the most preserved biological rhythms through evolution. It is composed by the states of wakefulness (W), non-rapid eye movement (NREM) sleep, and rapid eye movement (REM) sleep (1). These states are associated with different dynamical patterns of electric activity, which can be recorded accurately through the intracranial electroencephalogram, also known as electrocorticogram (ECoG).

In our previous work (2), we found that the ECoGs’ temporal complexity decreased from wakefulness to sleep; i.e., the repertoire of dynamical motifs was reduced when the animals fell asleep (Figure 1A, B and C). Interestingly, we observed this result in several cortical locations independent of its function (motor, olfactory, somatosensorial and visual), which suggested that the loss in temporal complexity was a global motif developed in the passage from W to sleep. Nevertheless, whether this result originated because a genuine change in brain dynamics happened or was a consequence of an artefactual measurement common to all recording electrodes, remained unanswered. It is important to consider this possibility because our previous recordings (2) employed a common reference in the cerebellum, which is in close proximity to the neck muscles and could be contaminated by the muscular activity.

**Figure 1.**
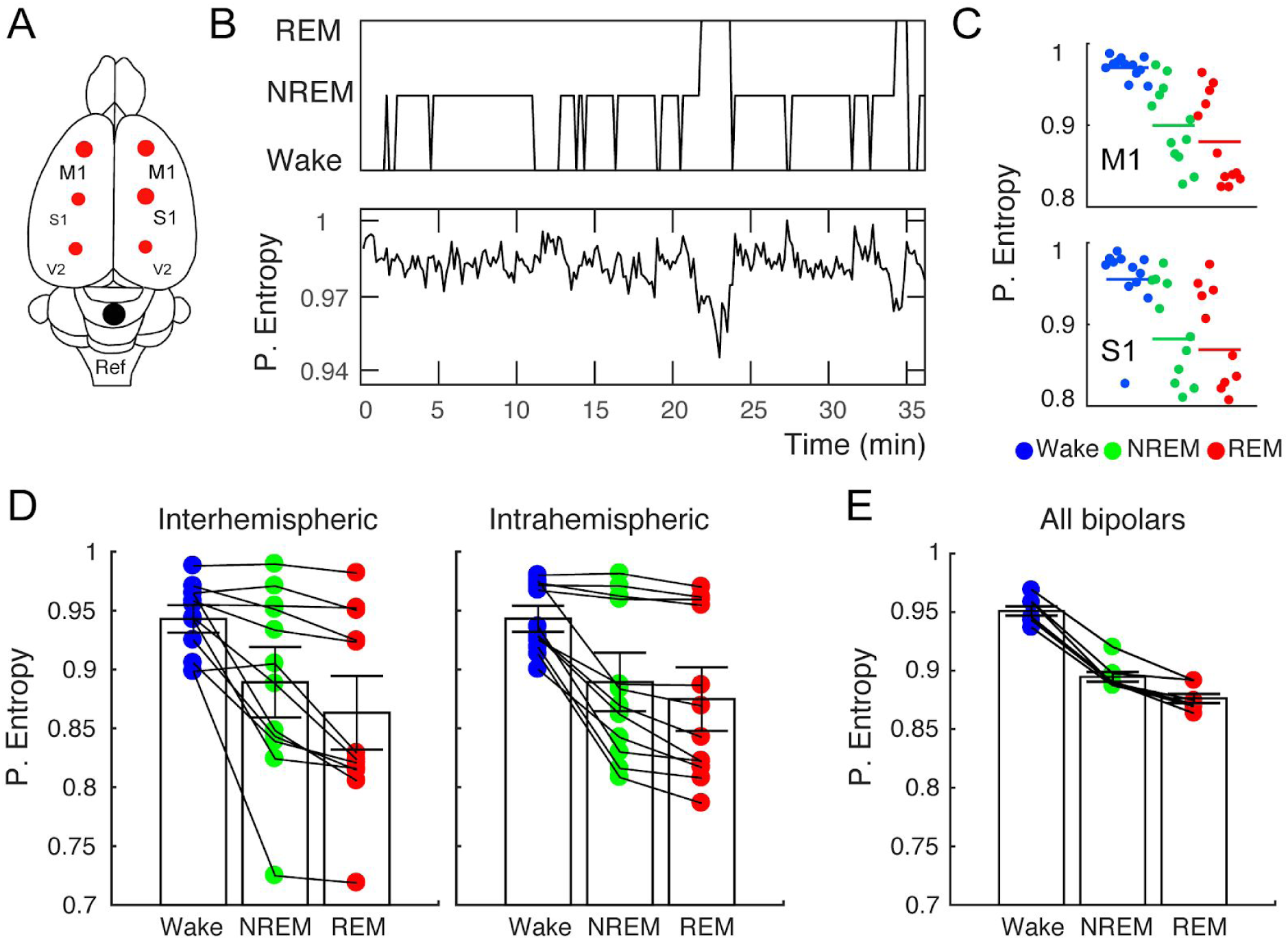
The ECoGs temporal complexity is independent of the electrode configuration. **A** Electrode localization in the rats cortex. The primary motor (M1; r and l, right and left) and right somatosensory (S1) cortex are shown together with the reference electrode placed above the cerebellum. **B** The hypnogram (top) from a representative animal is plotted simultaneously with the permutation entropy (bottom) from the M1r cortex as a function of time. **C** Scatter plots showing the time-average permutation entropy for each animal (12 rats) in each sleep state, blue W, green NREM and red REM. **D** The same scatter plots are now shown obtained from the bipolar configuration, interhemispheric (M1r-M1l) and intrahemispheric (M1r-S1r). **E** Permutation entropy decreases through sleep in all the bipolar configurations studied. Each dot depicts a bipolar electrode in each sleep and wake state (averaged from all the animals). 7 bipolars are plotted: M1r-M1l, M1r-S1r, M1l-S1l, S1r-S1l, S1r-V2r, S1l-V2l, V2r-V2l.

In order to discard this possibility, we re-referenced our data to obtain bipolar recordings, and then we measured their temporal complexity employing the same method as our previous work. Therefore, this approach removes the influence of the reference electrode and all common signals from our ECoGs, allowing us to investigate whether our previous results arise from a common background noise or were truly reflecting a global neural pattern which shifted from W to sleep.

## Material and Methods

In this report, we re-analyzed our previous data; therefore, the methods will be explained briefly and (2) should be consulted for a detailed description. We employed 12 Wistar adult rats maintained in a 12h light/dark cycle. All experimental procedures were conducted in agreement with the National Animal Care Law (No. 18611) and with the “Guide to the care and use of laboratory animals” (8th edition, National Academy Press, Washington DC, 2010). Furthermore, the Institutional Animal Care Committee (Comisión de Ética en el Uso de Animales) approved the experiments (Exp. No 070153-000332-16).

The animals were chronically implanted with electrodes to monitor the states of sleep and W. To record the ECoG, stainless steel screw electrodes were placed on the skull above motor (bilateral), somatosensory (bilateral), visual cortices (bilateral), the right olfactory bulb, and cerebellum, which was the reference electrode (see Fig. 1a and Table 2 in González et al. 2019). A neck bipolar electrode was employed to record the EMG. Experimental sessions were conducted during the light period, between 10 AM and 4 PM in a sound-attenuated chamber with Faraday shield. The recordings were performed through a rotating connector, to allow the rats to move freely within the recording box. Polysomnographic data were amplified x1000, acquired and stored in a computer using Dasy Lab Software employing 1024 as a sampling frequency and a 16 bits AD converter. The states of sleep and W were determined in 10 s epochs. W was defined as low voltage fast waves in the motor cortex, a strong theta rhythm (4-7 Hz) in the visual cortices, and relatively high EMG activity. NREM sleep was determined by the presence of high voltage slow cortical waves together with sleep spindles in motor, somatosensory, and visual cortices associated with a reduced EMG amplitude; while REM sleep as low voltage fast frontal waves, a regular theta rhythm in the visual cortex, and a silent EMG except for occasional twitches. An additional visual scoring was performed to discard artifacts and transitional states.

To assess the ECoGs temporal complexity, we employed the measure known as Permutation Entropy, which has been employed widely (3–7). This metric is robust to noise and it is computationally efficient. The permutation entropy is calculated as follows: we encoded the ECoGs time-series into ordinal patterns (OPs) by dividing the time-series into sequences of non-overlapping vectors (each containing 3 time stamps), and classifying them according to the relative magnitude of its elements. This transforms the graded EcoG time-series into a symbolic one, which can only contain up to six symbols maximum (factorial of the number of elements in a vector). Each symbol then represents a dynamical motif found in the ECoGs.

We note that small noise fluctuations are always introduced into the time-series in order to remove degeneracies; i.e., avoid the cases where, for example x(t) = x(t+1). After the symbolic time-series is obtained, the permutation entropy is calculated applying the Shannon Entropy (8) (SE = −∑*p*(*α*) log[*p*(*α*)]) to the probability distribution. Where *p*(*α*) is the probability (relative frequency of alpha in the symbolic time-series) of the *α* symbol. For the statistical analysis, we employed the repeated measures ANOVA and set p<0.05 to be considered significant.

## Results

In order to discard the contribution of extracranial noise to the complexity decrease during sleep, we generated bipolar recordings by subtracting two active electrodes. As our original data came as a differential recording to a common reference, the bipolar configuration eliminates the contribution of this electrode (9–11), in our case, the cerebellum. This is especially important because of the close proximity between our reference electrode and the neck muscles.

Figure 1D shows the results we obtained employing two anatomically relevant configurations: one an interhemispheric (M1r - M1l) and the other an intrahemispheric (M1r - S1r) combination. When we analyzed this new data, we found that the temporal complexity still decreased from W to NREM sleep and reached its lowest values during REM sleep. Furthermore, this result was observed in all the bipolar recordings analyzed (Figure 1D), irrespective of being inter or intrahemispheric combinations; notice that the complexity decrease during sleep is seen in every bipolar configuration employed (Figure 1E). When we investigated the origin of this complex activity, we found that the predominant temporal patterns were the monotonically increasing or decreasing ones (Figure 2A). This happened during W and was further overexpressed during sleep. It is worth noting that the frequency distribution of the bipolar ECoGs showed a power-law distribution which was steepened during the sleep states (Fig.2B), similar to our previous result found with monopolar electrodes (see Figure 2.B in (2)).

**Figure 2.**
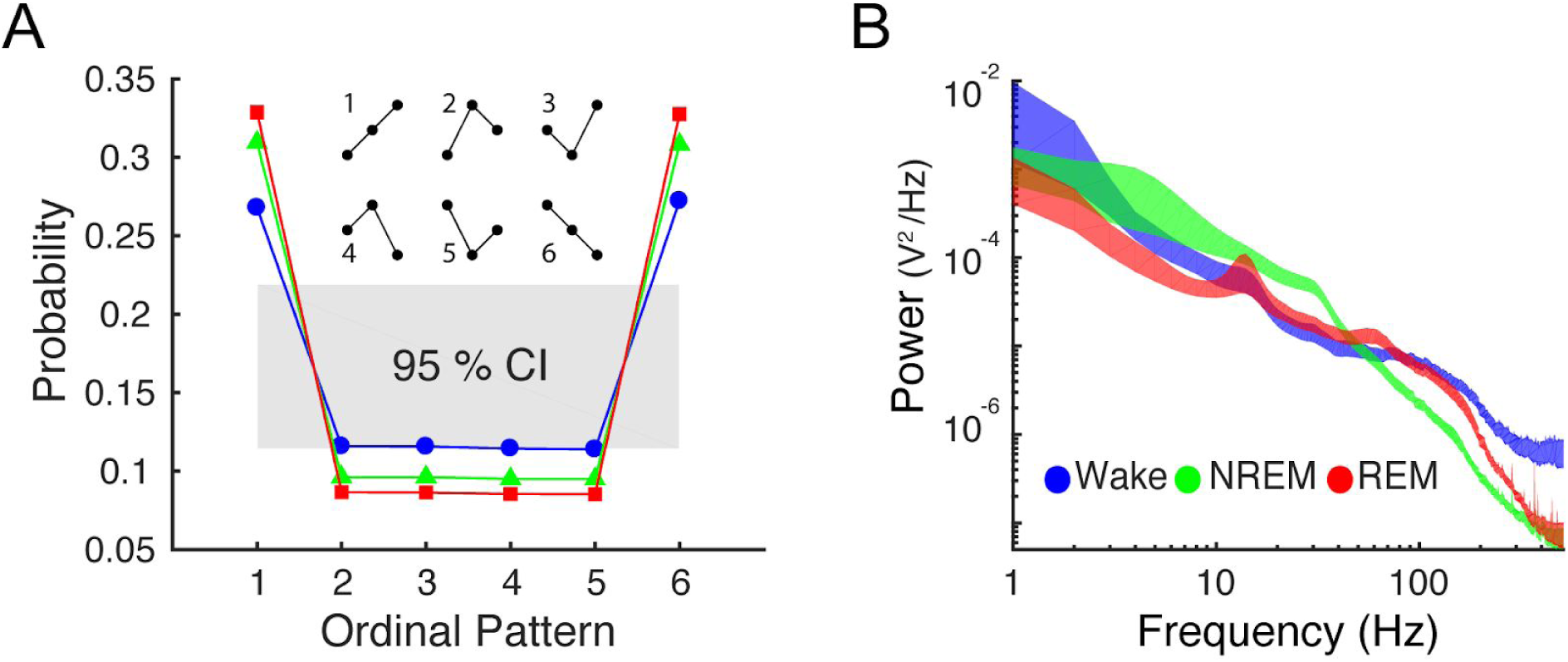
The dynamical characteristics of the ECoGs are preserved in the bipolar configuration. **A** Ordinal pattern probability distribution from the interhemispheric combination (M1r-M1l). The shaded area depicts the 95 percent confidence interval of the mean. The color code employed is the same as in panel **B. B** Average power spectral density (12 animals) during wakefulness and sleep, for the M1r-M1l bipolar configuration. The shaded areas depict the mean +/- the standard error.

## Discussion

In the present study we show that the loss in temporal complexity during sleep is not a consequence of a common noise entering through our reference electrode. This was evidenced by generating bipolar recordings, thus severely reducing the background noise common to all ECoG electrodes (9–12). This is particularly relevant because our reference electrode was closely located to the neck muscles and thus could be contaminated by the changes in muscle tone during sleep. In contrast, all bipolar recordings showed a significant complexity decrease as sleep progressed and reached its lowest values during REM. This means that our initial findings were independent on the electrode configuration employed (bipolar vs monopolar), and are less likely to simply reflect the changes in muscle activity during the sleep-wake cycle. Furthermore, the bipolar recordings retained a similar frequency and ordinal pattern distribution to what we had observed by the monopolar configuration, implying that these new changes in complexity arise from the same dynamic profile as in the monopolar case. Taken together, our results confirm that the electrocortical temporal complexity decreases from W to sleep, and this fact is not a consequence of a muscle artifact recorded through the reference electrode.

